# Greater cardiorespiratory fitness is associated with higher cerebral blood flow and lower oxygen extraction fraction in healthy older adults

**DOI:** 10.1101/2025.08.29.673075

**Authors:** Safa Sanami, Ali Rezaei, Stefanie A. Tremblay, Zacharie Potvin-Jutras, Dalia Sabra, Brittany Intzandt, Christine Gagnon, Amélie Mainville-Berthiaume, Lindsay Wright, Mathieu Gayda, Josep Iglesies-Grau, Anil Nigam, Louis Bherer, Claudine J. Gauthier

## Abstract

Aerobic exercise training promotes cardiovascular, brain and cognitive health. Regular exercise is associated with higher cardiorespiratory fitness, commonly assessed by peak oxygen uptake (VO_2peak_) during maximal effort testing. Higher cardiorespiratory fitness has been linked to preserved brain health, particularly higher grey matter volume and perfusion. The brain relies heavily on oxidative metabolism, yet the relationship between cardiorespiratory fitness and brain oxidative metabolism remains underexplored. This study investigated the association between VO_2peak_ and two key cerebral metabolic parameters: the cerebral metabolic rate of oxygen consumption (CMRO_2_) and oxygen extraction fraction (OEF), which represents the balance between cerebral blood flow (CBF) and CMRO_2_.

Thirty-seven healthy adults aged ≥50 underwent maximal cardiopulmonary exercise testing for VO_2peak_ assessment. Neuroimaging included dual calibrated functional MRI (dc-fMRI) and quantitative susceptibility mapping (QSM). Higher VO_2peak_ correlated positively with higher CBF across whole-brain grey matter but showed no relationship with CMRO_2_. Conversely, higher VO_2peak_ negatively correlated with lower OEF from both dc-fMRI and QSM. These findings suggest that greater cardiorespiratory fitness enhances cerebral perfusion without changing resting metabolic rate in healthy older adults, resulting in a reduced oxygen extraction. These results are consistent with exercise yielding improved vascular– metabolic coupling, which would reduce the likelihood of transient hypoxic episodes.

## Introduction

Cardiorespiratory fitness naturally declines with age but can be maintained or improved through regular aerobic exercise (1,2). While the gold standard for assessing cardiorespiratory fitness is maximal oxygen uptake (VO_2max_), due to practical limitations in older adults, peak oxygen uptake (VO_2peak_)—which closely corresponds to VO_2max_—is often used instead (3). Higher VO_2peak_ is consistently linked to lower all-cause mortality, improved cardiovascular health, and better cognitive and physical health outcomes in aging populations (4–8).

In the brain, regular aerobic exercise and higher cardiorespiratory fitness promote cognitive health, enhancing executive functions and memory in both younger and older adults (6,9,10). Structurally, higher cardiorespiratory fitness has been linked to greater cerebral volumes of grey and white matter, most notably in the hippocampus, but also in frontal and temporal regions (11,12). These structural enhancements likely underlie the improvements in cognitive performance observed with higher cardiorespiratory fitness (9). Functionally, cardiorespiratory fitness is thought to support cerebrovascular health by increasing cerebral blood flow (CBF), thereby improving oxygen and nutrient delivery to brain tissue (13–15).

Potentially underlying these beneficial effects, sustained exercise has been shown in both animal and human studies to promote angiogenesis, neurogenesis, and synaptic plasticity—particularly in the hippocampus (16,17). Moreover, exercise has been demonstrated in preclinical models to enhance mitochondrial function within neurons, contributing to more efficient brain energy metabolism and resilience against oxidative stress (18–21). Conversely, lower cardiorespiratory fitness has been linked to mitochondrial dysfunction (22), adversely impacting brain metabolism and overall brain health. Metabolic deficits and mitochondrial dysfunction may specifically manifest as a reduction in the cerebral metabolic rate of oxygen consumption (CMRO_2_), which represents the rate at which the brain consumes oxygen to support cellular metabolism (23,24). Another important biomarker sensitive to mitochondrial health is the oxygen extraction fraction (OEF), which reflects the balance between cerebral oxygen supply and utilization.

Improving cardiorespiratory fitness likely exerts beneficial effects on brain oxidative metabolism biomarkers, which can be non-invasively assessed using biomarkers such as CMRO_2_ and OEF. Investigating the relationship between VO_2peak_ and these metabolic markers is clinically relevant because they provide unique insights into brain function beyond traditional vascular metrics like CBF. For instance, elevated OEF could indicate a mismatch between oxygen delivery and cellular metabolic demands, suggesting an increased risk of transient ischemia (25). Similarly, reductions in CMRO_2_ might reflect compromised mitochondrial efficiency or neuronal loss (23,24). Thus, while higher fitness generally improves CBF, exploring CMRO_2_ and OEF specifically offers critical insights into the metabolic integrity and cellular health of the brain—dimensions not captured solely by vascular assessments. To date, only one study has investigated the relationship between whole-brain CMRO_2_ and VO_2peak_, reporting a negative association (26). However, no studies have yet examined the regional associations between cardiorespiratory fitness and CMRO_2_ or the global and regional associations with OEF. Understanding how these parameters vary with cardiorespiratory fitness in healthy adults may offer insight into the mechanisms linking fitness to brain health and whether exercise can be used to improve brain metabolic health.

In this study, we aimed to examine how VO_2peak_ relates to cerebral metabolic health in healthy adults aged 50 years and older. To do so, we assessed key vascular and metabolic biomarkers, including OEF, CBF, and CMRO_2_. OEF was measured using two complementary techniques: dual-calibrated functional MRI (dc-fMRI) and quantitative susceptibility mapping (QSM) to cross-validate our results, while CBF was measured using arterial spin labeling (ASL). We also derived estimates of CMRO_2_ by integrating these modalities. We then explored the associations between all brain parameters and VO_2peak_, offering novel insights into the links between fitness and brain metabolism.

## Methods

### Participants

Inclusion criteria in this study included being older than 50 years and fluent in either English or French to ensure informed consent and valid administration of neuropsychological assessments.

Exclusion criteria included a history of cardiovascular, neurological, psychiatric or respiratory disorder, thyroid disease, potential cognitive impairment (Mini Mental State Examination (MMSE) < 25), current tobacco use, high alcohol consumption (more than two drinks per day), contraindications to MRI (e.g., ferromagnetic implants, claustrophobia), having cardiometabolic risk factors known to affect brain vascular and metabolic health, including treated or untreated hypertension (BP > 140/90 mmHg) and diabetes (types 1 and 2). Participants were also excluded if they had undergone surgery under general anesthesia within the past 6 months, had malignant arrhythmias during exercise, arthritis or claudication, severe exercise intolerance, or excessive discomfort due to hypercapnia (> 5 on the Banzett dyspnea scale)(27).

A total of fifty-four (females; n=12) participants were recruited, of which 42 completed the study. Out of 12 participants that discontinued their participation, five participants did not complete the study due to interruptions during the COVID-19 pandemic, two participants were excluded due to claustrophobia during the MRI, two participants were excluded due to discomfort during the respiratory challenge necessary for the dc-fMRI and three participants chose not to participate for personal reasons (e.g., loss of interest, issues with scheduling). Of the 42 participants, 3 were unable to complete the full imaging protocol due to discomfort during the hypercapnia challenge—a controlled elevation of carbon dioxide levels used to assess OEF and CMRO_2_. These participants completed only the resting CBF acquisition and were therefore included in the CBF analysis but excluded from OEF and CMRO_2_ analyses. To ensure sufficient data quality, we excluded three participants with low-quality data based on the following criteria: pCASL temporal signal-to-noise ratio (tSNR) below 0.5 and excessive motion during the acquisition, defined as translational movements greater than 2mm, for a final sample of 37 participants with respiratory challenge. The study received approval from the Comité d’éthique de la recherche et du développement des nouvelles technologies (CÉRDNT) of the Montreal Heart Institute in accordance with the Declaration of Helsinki. All participants provided written informed consent prior to participation. Data collection was conducted at the Montreal Heart Institute.

Data was collected over three visits. Prior to the first visit, participants completed a medical history questionnaire to determine eligibility. Visit 1 included written informed consent, assessment of global cognitive function using the Mini-Mental State Examination (MMSE) (28). Additionally, participants underwent a 2-minute hypercapnia test to ensure their ability to tolerate the respiratory manipulation during the subsequent MRI session (with 2 participants being excluded at this stage). During the next visit, participants completed a stress test on a cycle ergometer to determine their peak oxygen consumption. The final visit was dedicated to the MRI acquisition. Demographic information for all participants is in Table 1.

**Table 1.**
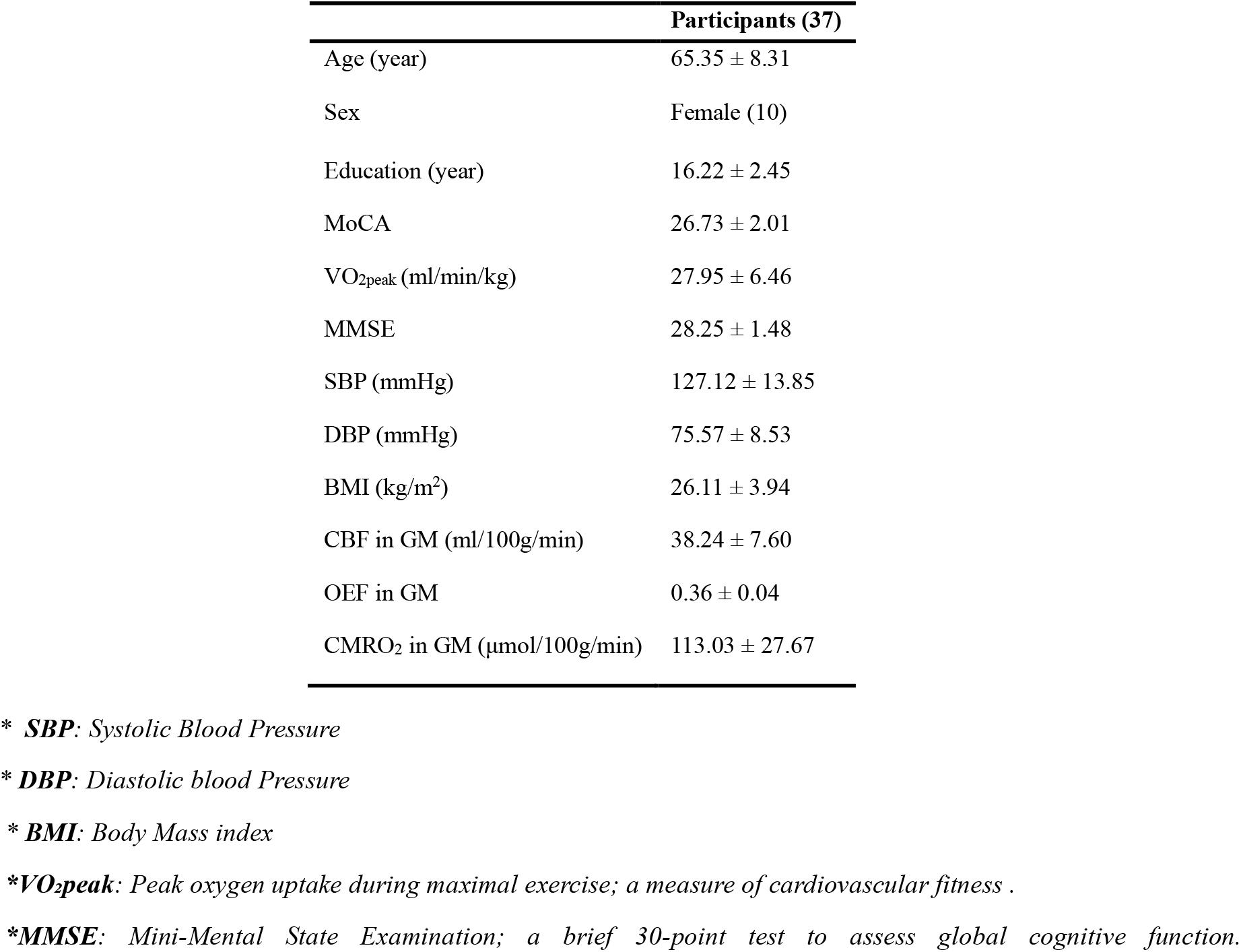
Demographic data on participants with respiratory data.

|  | Participants (37) |
| --- | --- |
| Age (year) | 65.35 ± 8.31 |
| Sex | Female (10) |
| Education (year) | 16.22 ± 2.45 |
| MoCA | 26.73 ± 2.01 |
| VO<sub>2peak</sub> (ml/min/kg) | 27.95 ± 6.46 |
| MMSE | 28.25 ± 1.48 |
| SBP (mmHg) | 127.12 ± 13.85 |
| DBP (mmHg) | 75.57 ± 8.53 |
| BMI (kg/m<sup>2</sup>) | 26.11 ± 3.94 |
| CBF in GM (ml/100g/min) | 38.24 ± 7.60 |
| OEF in GM | 0.36 ± 0.04 |
| CMRO<sub>2</sub> in GM (μmol/100g/min) | 113.03 ± 27.67 |

\* **SBP**: Systolic Blood Pressure
\* **DBP**: Diastolic blood Pressure
\* **BMI**: Body Mass index
\***VO<sub>2peak</sub>**: Peak oxygen uptake during maximal exercise; a measure of cardiovascular fitness .
\***MMSE**: Mini-Mental State Examination; a brief 30-point test to assess global cognitive function.

### Maximal cardiopulmonary exercise test (CPET)

Incremental CPET to volitional exhaustion was performed on a cycle ergometer (E100, Cosmed, Italy) according to the latest recommendations and as previously published (29,30). A 3-minute warm-up phase at an initial workload of 20 W was followed by an incremental exercise test, with increases of 10 to 20 W per minute depending on the participant’s physical capacity, performed at a pedaling cadence between 60 and 80 rpm. The recovery phase consisted of 2 minutes of active recovery at 20 W at pedalling speed between 50 and 60 rpm, followed by 3 minutes of passive recovery, during which participants remained seated quietly on the ergometer. Gas exchange parameters were continuously measured at rest, during exercise, and during recovery using a metabolic system (Cosmed Quark, Cosmed, Italy), capturing on a breath-by-breath basis and then averaged in 10 second increments as recently published (29). There was continuous Electrocardiogram (ECG) monitoring before, during the test and in the recovery (T12x, Cosmed, Italy). Diastolic (DBP) and systolic (SBP) blood pressures (Tango M2, Suntech, USA) and ratings of perceived exertion (RPE) were measured at rest and every 3 minutes throughout the test. The protocol interrupted if one of the following happened: (1) inability to maintain the required cadence of 60 rpm despite verbal encouragement, (2) clinical indications requiring interruption of the test, such as volitional exhaustion, (3) abnormal ECG findings or angina and, (4) excessive systolic blood pressure response (210 mmHg for men, 190 mmHg for women (31)). The highest VO_2_ value reached during the exercise phase, averaged over a 10-second interval, was considered VO_2peak_ and expressed in mL/min/kg of lean body mass. Lean body mass was estimated using a body composition analyzer (TANITA), which also provided measurements of fat mass.

### MRI acquisition

Data were collected using a 3T Skyra MRI system with a 32-channel head coil. The acquisition protocol included structural MRI, dual-echo pseudo-continuous Arterial Spin Labeling (pCASL) for simultaneous acquisition of perfusion and Bold Oxygen Level Dependent (BOLD) signals and a blood magnetization map (M0) for perfusion quantification. pCASL data were acquired with a voxel resolution of 3.43 × 3.4375 × 7 mm, TR/TE1/TE2/alpha: 4000/10/30 ms/90°, labeling duration of 1517 ms with a post-labeling delay (PLD) of 1300 ms. The M0 acquisition had identical parameters but with a TR of 10 seconds, to ensure a fully recovered magnetization. A 3D multi-coil multi-echo gradient echo (ME-GRE) (TR/TE1/TE2/TE3/TE4/flip angle = 20/6.92/13.45/19.28/26.51 ms/9°, 0.7 × 0.7 × 1.4 mm^3^ voxel size) phase and magnitude data were acquired for all the coils separately for the reconstruction of QSM maps. Flow compensation of the first TE was completed by nulling the gradient moment to ensure minimizing flow artifact effects for the venous mapping (32).

As the MRI data were acquired within two sub-sessions to allow the participant to take off the mask for the rest of the data acquisition, two T1-weighted acquisitions were collected to ensure registration accuracy of the pCASL slab. One lower resolution acquisition was performed right before pCASL while the participant had the mask on, and a higher resolution acquisition was performed during the second sub-session, without the mask. The low resolution T1-weighted acquisition was acquired using a magnetization prepared rapid gradient echo (MPRAGE) sequence with TR/TE/Flip angle = 15 ms/3.81 ms/25° with a 1.5 mm isotropic resolution. The high-resolution T1-weighted structural images were also acquired with an MPRAGE sequence, with TR/TE/Flip angle = 2300 ms/2.32 ms/8° with a 0.9 mm isotropic resolution.

### Respiratory manipulation

The RespirAct™ system (RespirAct_TM_, Thornhill Research, Toronto, Canada) was used during the breathing manipulation to target specific end-tidal partial pressures of CO_2_ and O_2_. Three conditions were sequentially targeted for 2 minutes, preceded and followed by 2 minutes of room air inhalation: a hypercapnic condition (targeting 5 mmHg above baseline for end-tidal CO_2_), an isocapnic hyperoxic condition (targeting 150 mmHg above baseline for end-tidal O_2_), and a combined hyperoxia and hypercapnia condition (targeting 5 mmHg above baseline for end-tidal CO_2_ and 150 mmHg for end-tidal O_2_). A rebreathing face mask was used to deliver gases to the participants. The system delivered gas at a flow rate of 20 L/min, while the concentrations of exhaled CO_2_ and O_2_ were monitored. On their first study visit, a familiarization session was done with a 2-minute hypercapnic manipulation to ensure participant comfort during the MRI. Breathing discomfort was assessed using the Banzett dyspnea scale and those with a score >5 were not invited to continue in the study (n = 2) (27).

### Respiratory data analysis

CO_2_ and O_2_ were sampled continuously throughout the breathing manipulation by the RespirAct, which also generates a time series of end-tidal CO_2_ and O2 partial pressures. The end-tidal partial pressure is used as a proxy for arterial gas concentrations (33). To combine these values with the MRI data, a MATLAB script was employed to smooth the data, remove outliers and resize the data to match the durations of the BOLD and ASL signals.

### Imaging data processing

#### T1 processing

All structural images underwent preprocessing with the Brain Extraction Tool (BET) in FSL to remove the skull. Subsequently, FSL’s FAST was employed to segment the structural images into Grey Matter (GM), White Matter (WM), and Cerebrospinal Fluid (CSF).

### Preprocessing of BOLD and pCASL

Both ASL and BOLD datasets were corrected for motion using mcflirt from FSL. Then, a mask was created using the motion corrected BOLD images and BET, and applied to both echoes to remove the scalp. The first echo of the dual-echo pCASL acquisition was used to compute the perfusion-weighted time-series, from which CBF was estimated using surround subtraction. Volumes that contained voxels with intensity values outside of 3 standard deviations from the mean were flagged. Then, volumes with more than 50% of voxels identified as outliers were excluded from further analysis, to ensure adequate tSNR (34). The M_0_ image underwent skull stripping via BET and was utilized for calibrating the perfusion image. The second echo was used to extract the BOLD-weighted signal via surround addition, reflecting changes in blood oxygenation.

### CBF maps using pCASL data

Perfusion was quantified from the preprocessed ASL time-series using FSL’s BASIL for kinetic modeling combined with the M_0_ acquisition (34). Finally, partial volume correction was applied using a weighted average taking into account tissue contributions using the segmentation results of GM, WM, and CSF from the T1-weighted data as in (35).

### OEF & CMRO_2_ maps using calibrated fMRI

OEF was estimated using the general calibration model (GCM), based on the deoxyhemoglobin dilution model of the BOLD signal (36). Following OEF estimation, CMRO_2_ was calculated using CBF maps and arterial oxygen content using Fick’s principle, as described in (Gauthier & Hoge, 2012)

### QSM data reconstruction

To reconstruct the QSM maps, the magnitude data of the coils were combined by the square root of the sum of squares of each coil data for each TE (37). The phase data of multi-coil ME-GRE data was combined and unwrapped using the ROMEO toolbox to ensure spatiotemporal coherence and temporal stability (38). Then, the QSM (X) maps were reconstructed using the first echo data with Total-Generalized Variation (TGV) QSM reconstruction method (37). The TGV algorithm minimizes the noise propagation through consecutive QSM processing steps and preserving edges between different tissue types, especially the edges between veins and surrounding tissue. The QSM maps were zero-referenced by the mean values of the CSF in ventricles (39). The veins were extracted using a multi-scale recursive ridge filtering method (40) from the QSM data reconstructed from the first TE data which was the only echo time with flow compensation. OEF is directly related to the concentration of paramagnetic deoxyhemoglobin present in veins using the following formula:

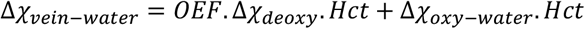

where the Δχ_*deoxy*_= 0.27 ppm (cgs) is the susceptibility shift (per unit hematocrit) between the fully oxygenated and fully deoxygenated red blood cells, and Δχ_*oxy*−*water*_= -0.03 ppm (cgs) is the susceptibility shift of oxygenated red blood cells. Hematocrit (Hct) values were considered 40% (41).

### Registration of QSM and calibrated fMRI to MNI space

The pCASL images were registered to MNI space in three separate steps. First, a rigid body registration with nearest neighbor interpolation was conducted using Advanced Normalization Tools (ANTs) (42) to align the pCASL mean images with the low-resolution (1.5 mm) T1-weighted image. Next, the low-resolution T1-weighted images were registered to the high-resolution (0.9 mm) T1 space using a multistage approach that combined rigid, affine, and nonlinear (SyN) transformations with a multi-resolution strategy. In the final step, the high-resolution T1 images were registered to MNI space using the same transformation method applied for the registration from low to high resolution. Finally, the pCASL images were registered to MNI space by applying the three transformation matrices obtained from the previous registration steps, utilizing 3 warp images. The QSM images were transformed to the high-resolution T1 space and then to the MNI space using the same methods.

After registering all images to MNI space, we extracted regional CBF, OEF, and CMRO_2_ from the pCASL data using the LPBA40 atlas (43). Mean CBF values were calculated for each region; however, due to incomplete coverage and low signal quality in some areas, only 44 of the 56 atlas regions were retained for analysis. Regions such as the occipital, lingual, and fusiform cortices were excluded for these reasons. We also included overall grey matter CBF, resulting in a total of 45 regions analyzed. For the QSM data, because the number of veins detected require larger regions to be reliable, mean OEF values were extracted from veins in frontal, temporal, parietal, and occipital lobes using the MNI structural atlas.

### Statistical analysis

Normality of regional data distributions was assessed using the Shapiro–Wilk test. For each biomarker— CBF from calibrated fMRI, OEF from both calibrated fMRI and QSM, and CMRO_2_ from calibrated fMRI— we performed linear regression analyses with VO_2peak_ as the predictor, while controlling for age and sex as covariates. To account for multiple comparisons, we applied false discovery rate (FDR) correction to the resulting p-values. To assess agreement between the two OEF estimation methods, a Bland‐Altman analysis was used, which evaluates the mean difference and limits of agreement between paired measurements. This approach quantitatively characterizes systematic bias and variability between whole-brain OEF derived from dual-calibrated fMRI and OEF values from QSM across subjects. A priori p-values were set to ≤ 0.05.

### Comparison of Whole-Brain OEF between dc-fMRI and QSM

To evaluate the agreement between methods, whole-brain OEF estimates obtained from dc-fMRI were compared with venous OEF derived from QSM. This comparison was conducted using Bland-Altman analysis. Details of the analysis and corresponding results are provided in the supplementary data and Supplementary Figure 1.

## Results

All regional data distributions passed the Shapiro–Wilk test for normality (p > 0.05), indicating that the normality assumption was met.

### CBF and VO_2peak_

We found widespread positive associations between CBF and VO_2peak_ in whole gray matter, with regional effects in bilateral frontal, temporal, as well as subcortical structures (Figure 1A). A full list of regions is provided in Table 2.

**Figure 1.**
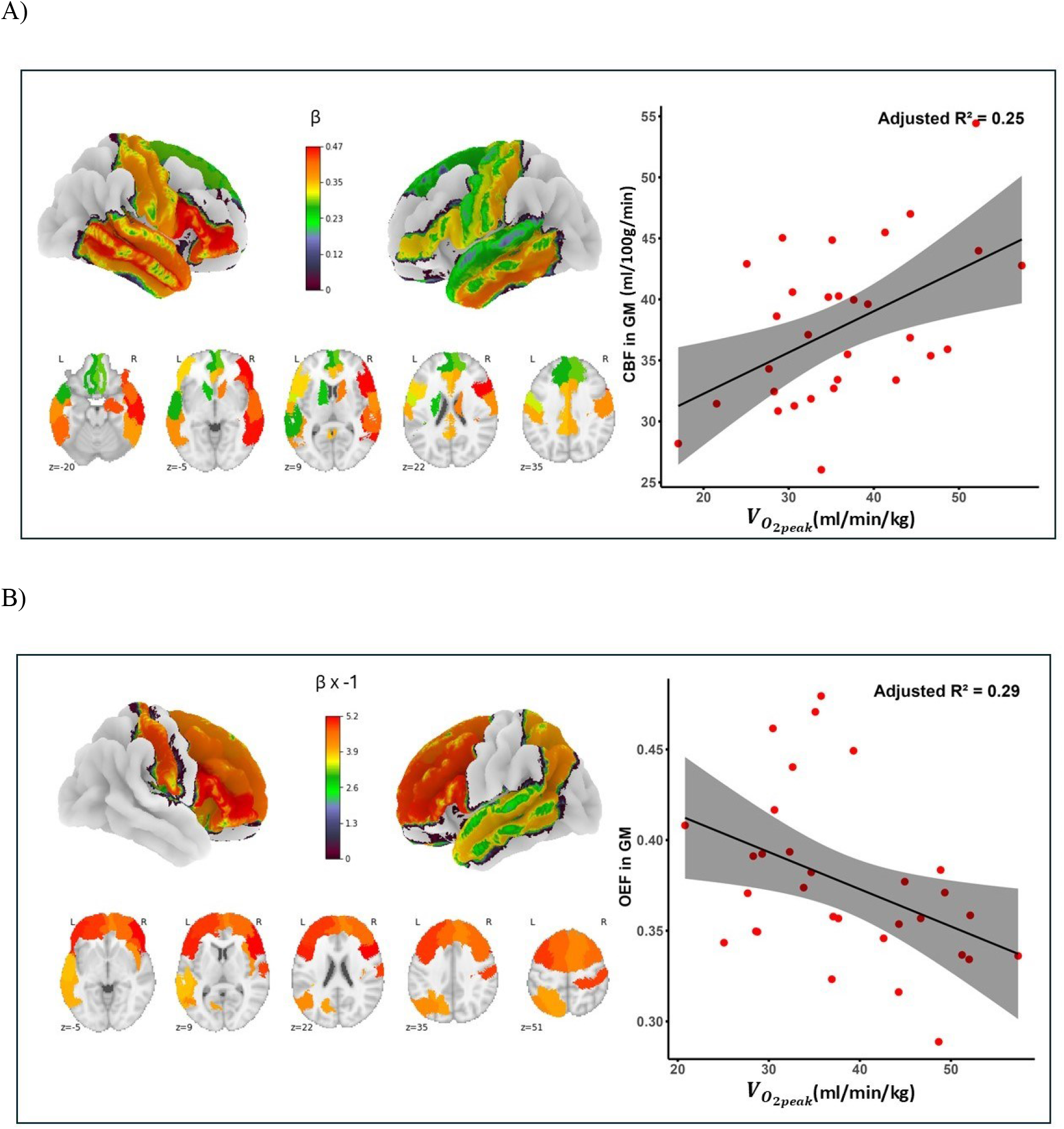
Correlations between VO_2peak_ and CBF and between VO_2peak_ and OEF. A) Positive association between VO_2peak_ and CBF measured using pCASL. β coefficient maps (multiplied by 10^-3^ for ease of visualization) (left) are overlaid on 3D brain surface, and corresponding scatterplots (right) show linear relationship between CBF in GM and VO_2peak_, with 95% confidence intervals shaded. B) Negative association between VO_2peak_ and OEF estimated from dc-fMRI. β coefficient maps (multiplied by -1 for ease of visualization) (left) are overlaid on 3D brain surface, and corresponding scatterplots (right) show linear relationship between OEF in GM and VO_2peak_, with 95% confidence intervals shaded.

**Table 2.**
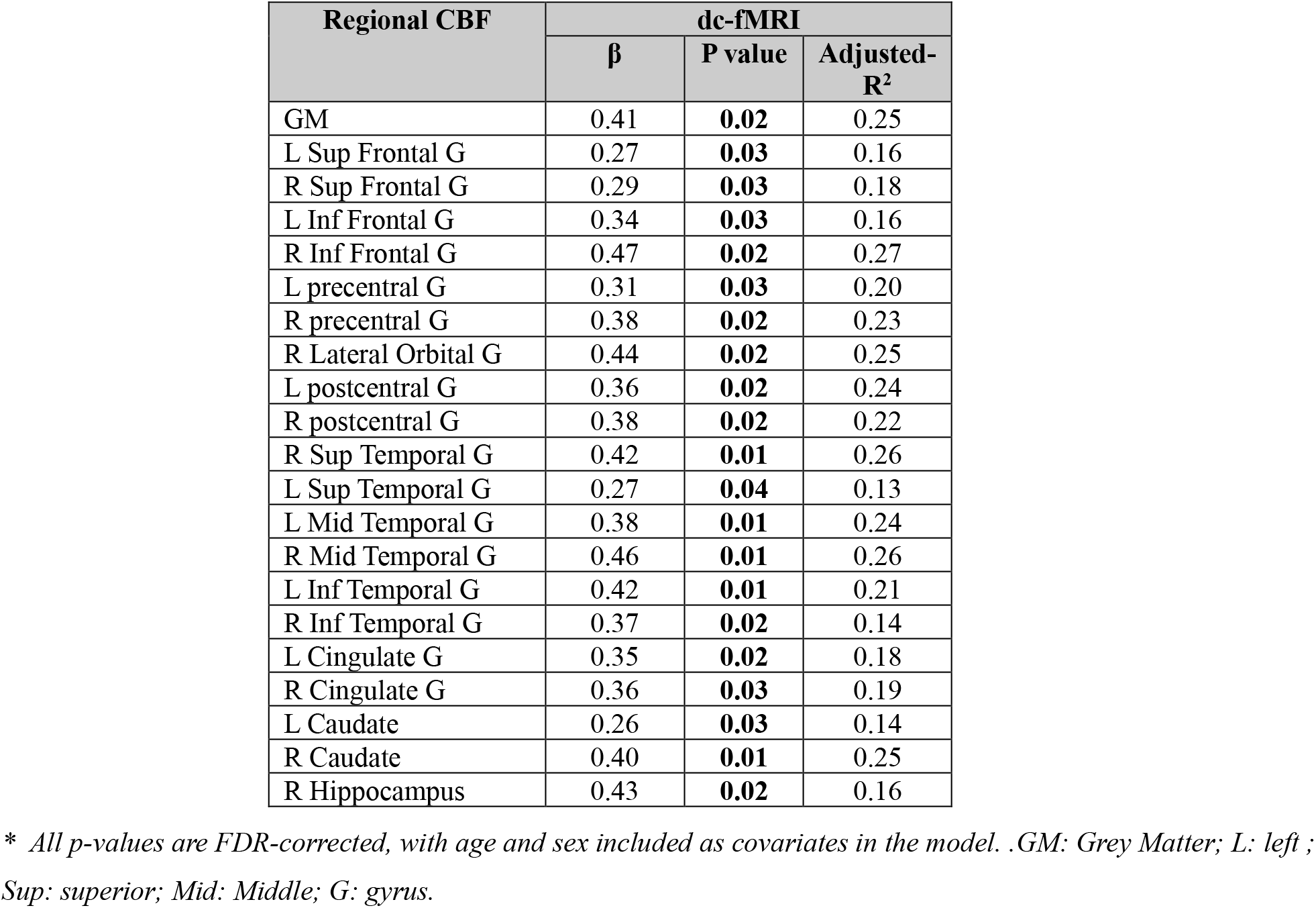
Brain regions showing significant associations between CBF and VO_2peak_.

### CMRO_2_ and VO_2peak_

We observed no significant associations between CMRO_2_ and VO_2peak_ in any brain region (p > 0.05).

### OEF and VO_2peak,_

In our dc-fMRI data, we observed a negative association between OEF and VO_2peak_ across multiple brain regions (p < 0.05), including gray matter, and several frontal, parietal, temporal, and insular areas (Figure 1B), as detailed in Table 3. Similarly, our linear regression analysis on the QSM-derived data confirmed this negative association at the whole-brain level as well as in the frontal, temporal, occipital, and parietal lobes (Figure 2), as shown in Table 3.

**Table 3.** Brain regions showing a significant negative association between VO_2peak_ and OEF in both dc-fMRI and QSM techniques.

| Regional OEF | dc-fMRI |  |  |
| --- | --- | --- | --- |
| | $\beta$ | P value | Adjusted-R<sup>2</sup> |
| GM | -3.60 | **0.02** | 0.29 |
| L Sup Frontal G | -4.62 | **0.02** | 0.24 |
| R Sup Frontal G | -4.29 | **0.02** | 0.21 |
| L Mid Frontal G | -4.86 | **0.02** | 0.25 |
| R Mid Frontal G | -4.60 | **0.02** | 0.27 |
| L Inf Frontal G | -4.99 | **0.02** | 0.31 |
| R Inf Frontal G | -5.17 | **0.01** | 0.35 |
| R Lateral Orbital G | -4.74 | **0.02** | 0.25 |
| R postcentral G | -4.84 | **0.03** | 0.19 |
| L Sup Parietal G | -4.03 | **0.02** | 0.21 |
| L Angular G | -4.21 | **0.02** | 0.21 |
| L Sup Temporal G | -3.87 | **0.02** | 0.25 |
| L Mid Temporal G | -3.92 | **0.02** | 0.28 |
| R Insula | -4.17 | **0.02** | 0.21 |
|  | QSM |  |  |
| | $\beta$ | P value | Adjusted-R<sup>2</sup> |
| Whole brain | -0.001 | **0.02** | 0.20 |
| Frontal | -0.001 | **0.02** | 0.21 |
| Parietal | -0.001 | **0.03** | 0.12 |
| Temporal | -0.002 | **0.01** | 0.27 |
| Occipital | -0.002 | **0.02** | 0.18 |

\* $\beta$ values in dc-fMRI methods should be multiplied by $10^{-3}$
Linear regression results showing the association between $VO_{2peak}$ and OEF in dc-fMRI and QSM-derived data across selected brain regions. All models include age and sex as covariates. Reported p-values have been FDR-corrected to account for multiple comparisons. GM: Grey Matter; L: left; Sup: superior; Mid: Middle; G: gyrus.

**Figure. 2.**
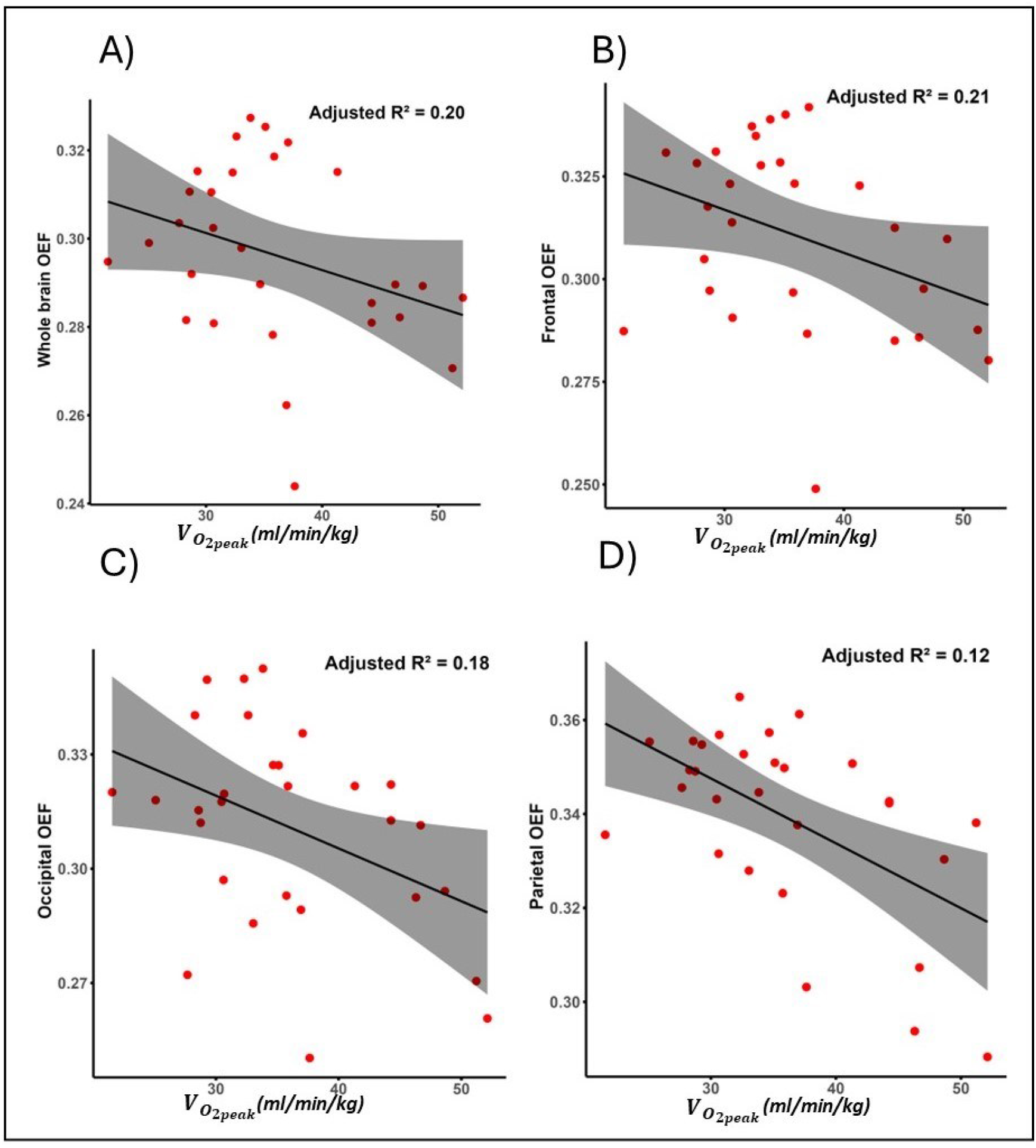
Associations between VO_2peak_ and regional venous OEF derived from QSM. The scatter plots illustrate significant inverse relationships between VO_2peak_ and OEF in the whole brain (A), frontal lobe (B), occipital lobe (C), and parietal lobe (D). Shaded areas represent 95% confidence intervals.

## Discussion

In this study, we investigated the relationship between VO_2peak_ and key physiological brain biomarkers, including vascular (CBF) and metabolic (OEF and CMRO_2_) measures, in healthy older participants. To our knowledge, this is the first study to examine the association between VO_2peak_ and regional brain metabolism biomarkers. Our results revealed a positive relationship between VO_2peak_ and CBF across the whole grey matter, as well as in the frontal, temporal, and subcortical regions, including the hippocampus and caudate. In contrast, we found no significant relationship between VO_2peak_ and CMRO_2_ in any brain region. Finally, negative associations between VO_2peak_ and OEF measured using dc-fMRI were observed in the whole brain, grey matter, frontal and temporal lobes, and insula. With QSM, the negative associations extended to the entire brain and all major lobes, including the frontal, temporal, occipital, and parietal regions. This study demonstrates for the first time that, in healthy adults over 50, cardiorespiratory fitness is linked with higher oxygen availability, likely contributing to protecting the brain against transient ischemic episodes.

Our results of a positive relationship between VO_2peak_ and CBF is in agreement with some studies, which have reported positive associations in grey matter regions, including the hippocampus and temporal and occipital lobes (13,44,45). However, the direction of association remains inconsistent in the literature, with some studies having reported negative or null associations, particularly among cohorts with high baseline fitness (10,16,46,47). These mixed results may reflect variations in participants’ fitness levels, the presence of vascular pathology and risk factors, and different methodological approaches across studies. Our findings align with the exercise-related vascular adaptations, such as increased capillary density—observed in animal models (48,49)—, as well as exercise-induced upregulation of growth factors like brain-derived neurotrophic factor (BDNF) and vascular endothelial growth factor (VEGF), which are known to promote angiogenesis (50,51). However, future studies in larger cohorts are needed to disentangle the parameters that could affect this relationship, including sex, age, fitness level, and the presence of other factors known to affect CBF such as hypertension.

Conversely, we observed no significant relationship between CMRO_2_ and VO_2peak_, suggesting that cerebral metabolic demands remain relatively stable across fitness levels in healthy older adults. While there is a paucity of studies in the literature to contextualize our findings, functional imaging studies using techniques such as functional near-infrared spectroscopy (fNIRS) have indicated that higher fitness may positively influence brain oxygenation during cognitive tasks or exercise, primarily among older adults (52,53). Interestingly, one previous study reported a negative relationship between global CMRO_2_ and VO_2peak_ (26). However, participants in that study were generally older and were not selected to be free of hypertension and other vascular risk factors. Furthermore, higher global CMRO_2_ in that study was also related to fatigability and multi-morbidity, suggesting that there may be a nonlinear relationship whereby CMRO_2_ may increase as a compensatory mechanism in some conditions. This is for example consistent with findings relating higher fatigability to higher CMRO_2_ in multiple sclerosis (54). While we did not collect fatigability information in this study, future studies should attempt to disentangle this potentially nonlinear relationship between VO_2peak_ and CMRO_2_ depending on health status and fatigability.

Our study breaks new ground in documenting for the first time the relationship between OEF and VO_2peak_. We observed a negative relationship between VO_2peak_ and cerebral OEF, suggesting that healthy individuals with higher cardiovascular fitness achieve greater cerebral oxygen delivery without increasing oxygen utilization, consistent with the absence of a corresponding relationship with CMRO_2_. To our knowledge, this is the first study to demonstrate this relationship, which we confirmed using two independent imaging techniques. This lower OEF, combined with unchanged CMRO_2_ and elevated CBF is consistent with a greater delivery of O_2_ to tissue, but since the same absolute amount of O_2_ is extracted from arterial blood, more O_2_ remains in venous blood. This improved vascular–metabolic coupling may be beneficial by ensuring that oxygen supply remains adequate even when the need for O_2_ transiently increases, thereby lowering the likelihood of transient hypoxic episodes. Clinically, elevated OEF has been associated with tissue at greater risk of further ischemic damage after stroke (25), highlighting the relevance of optimal oxygen delivery for maintaining brain health.

Exploring the relationships between VO_2peak_ and these cerebral vascular and metabolic biomarkers is clinically valuable, as changes in OEF and CMRO_2_ may offer sensitive indicators of mitochondrial dysfunction. Mitochondrial impairment is characterized by a reduced ability to utilize oxygen (55), leading to concurrent reductions in CMRO_2_ and OEF if cerebral blood flow remains relatively stable. Conversely, elevated OEF values typically reflect vascular impairment, such as increased vessel stiffness or reduced endothelial function, often observed with aging or cardiovascular risk factors (56,57). Thus, understanding whether higher aerobic fitness associates with alterations in these biomarkers in healthy adults could shed light on the underlying mechanisms through which exercise interventions confer brain health benefits. These results could inform targeted preventive strategies in populations vulnerable to metabolic and cerebrovascular dysfunction.

By applying both calibrated fMRI and QSM, our study demonstrates consistent associations between VO_2peak_ and OEF, reinforcing the robustness of our findings. Importantly, we found that QSM‐derived OEF values are systematically lower by ∼15% compared to dc-fMRI, with no evidence of proportional bias across the range. These lower values could be due to the greater sensitivity of QSM to partial volume effects, as well as different contributions of GM and WM since it is unknown which tissue type drains into the veins detected by QSM. Lack of proportional bias and relatively narrow limits shows that QSM is reliably tracking the same physiological effect as dc-fMRI OEF. Hence, QSM emerged as a practical alternative in settings where calibrated fMRI may be difficult to implement. The latter, while powerful, requires respiratory manipulations which can be uncomfortable and rely on expensive specialized equipment, as well as prolonged scan times, which can pose challenges—especially for older adults or clinical populations. In contrast, QSM is a non-invasive, faster, and more comfortable alternative, making it particularly well-suited for broader clinical applications. As an initial exploration of fitness-related brain metabolic patterns, our study lays the groundwork for future investigations in patient populations, where understanding these relationships could inform more personalized and targeted interventions.

## Limitations

The primary limitation of this study is the small sample size, which resulted from the difficulty of finding subjects who met our strict inclusion and exclusion criteria, and possibly from participant attrition due to discomfort experienced during the hypercapnia challenge. This limitation may reduce the generalizability of our findings. Future studies with larger cohorts and a wider range of health status are needed to further support our results.

ASL suffers inherently from a low tSNR, which can be exacerbated by prolonged arterial transit time in populations with vascular risk. A single PLD, as employed here, therefore makes ASL vulnerable to underestimation of CBF in individuals with delayed transit time. Future studies should use multi-PLD sequences to improve CBF quantification, and use background suppression to improve SNR. Background suppression was not used here to maintain BOLD SNR in our dual-echo acquisition for OEF and CMRO_2_ quantification.

The accuracy of venous OEF values from QSM depends critically on the quality of the vein segmentation and on partial-volume effects—particularly in veins whose diameter is significantly smaller than the image resolution—which tend to cause systematic underestimation. Thus, future studies should develop vein segmentation techniques that are more sensitive to the veins with smaller diameter will help to decrease the effect of partial volumes. Furthermore, future studies should use susceptibility reconstruction techniques that better delineate between veins and surrounding tissue, such as X-separation techniques, which facilitates vein segmentation from paramagnetic and diamagnetic susceptibility maps.

## Conclusion

This study provides new evidence that higher cardiorespiratory fitness, measured by VO_2peak_, supports better oxygen delivery to the brain in healthy adults. Participants with greater fitness had increased blood flow to the brain, enabling enhanced oxygen availability without raising the brain’s overall metabolic demand or the need for greater oxygen extraction. Additionally, we showed that QSM, an imaging method not requiring respiratory challenges, provides similar estimates of brain oxygen use compared to traditional calibrated MRI, making it a useful alternative, especially in populations who cannot comfortably tolerate breathing challenges. Together, these results offer new insights into the neurovascular effects of fitness and provide a foundation for future studies exploring these relationships in aging and clinical populations, where targeted interventions may benefit from individualized assessments of brain oxygenation and perfusion.

## Acknowledgments

We would like to thank everyone who contributed to this project: Paule Samson, Thomas Vincent, Julie Lalongé, Hakima Benhalima, Milla Shakleva, Victoria D’Amours, Agathe Godet, Stephanie Beram, Roni Zaks, Robert Hovey, Alexandre Bailey, Catherina Medeiros, Amélie Mainville-Berthiaume and Zineb Rouabah. Thank you also to the laboratories of Dr Louis Bherer and Dr Mathieu Gayda. Lastly, we would like to acknowledge our research participants without whom none of this would have been possible.

## Author Contributions

Safa Sanami and Ali Rezaei contributed equally to this work, performing the main analysis, data acquisition, and manuscript writing.

Stefanie A. Tremblay, Zacharie Potvin-Jutras, and Dalia Sabra contributed to data acquisition, manuscript review, and editing.

Brittany Intzandt, Christine Gagnon, Amélie Mainville-Berthiaume, Lindsay Wright, Mathieu Gayda, Josep Iglesies-Grau, and Anil Nigam contributed to manuscript review and editing.

Louis Bherer and Claudine J. Gauthier contributed significantly to conceptualization, main methodology, manuscript editing, and review.

## Disclosure

None.

The author(s) declared no potential conflicts of interest with respect to the research, authorship, and/or publication of this article.

## Funding

This work was supported by funding awarded to Claudine J. Gauthier from the Natural Sciences and Engineering Research Council of Canada (NSERC Discovery Grant: RGPIN-2015-04665; 2024-06455), Fonds de recherche du Québec (FRQ 5232), the Heart and Stroke Foundation of Canada (G-17-0018336), the Heart and Stroke Foundation New Investigator Award, the Henry J.M. Barnett Scholarship, the Michal and Renata Hornstein Chair in Cardiovascular Imaging and the Mirella and Lino Saputo research chair in cardiovascular health and the prevention of cognitive decline. Additional support was provided by the Vascular Training Platform (VAST) (to Ali Rezaei), the Canadian Institutes of Health Research (FRN: 175862, to Stefanie A. Tremblay), the Heart and Stroke Foundation of Canada and Brain Canada (to Zacharie Potvin-Jutras), and the Alzheimer Society Research Program Postdoctoral Award (to Brittany Intzandt.).

## Notes

### Competing Interest Statement

The authors have declared no competing interest.

