## Supplementary material for "Greater cardiorespiratory fitness is associated with higher cerebral blood flow and lower oxygen extraction fraction in healthy older adults": Figure S1

*dc-fMRI OEF vs. QSM venous OEF*

For the whole brain OEF, dc-fMRI yielded systematically higher OEF than QSM. Mean ± SD values were 0.35 ± 0.05 for dc-fMRI and 0.3 ± 0.02 for QSM. Bland-Altman analysis showed a bias of +0.05 OEF units ($p < 0.001$), with 95 % limits of agreement ranging from –0.04 to +0.13 (Figure S1). Although most observations fell within these limits, the spread widened compared with the cortical average, and the negative bias (~15 % of the dc-fMRI mean) indicates that QSM tends to underestimate the whole brain OEF relative to dc-fMRI.

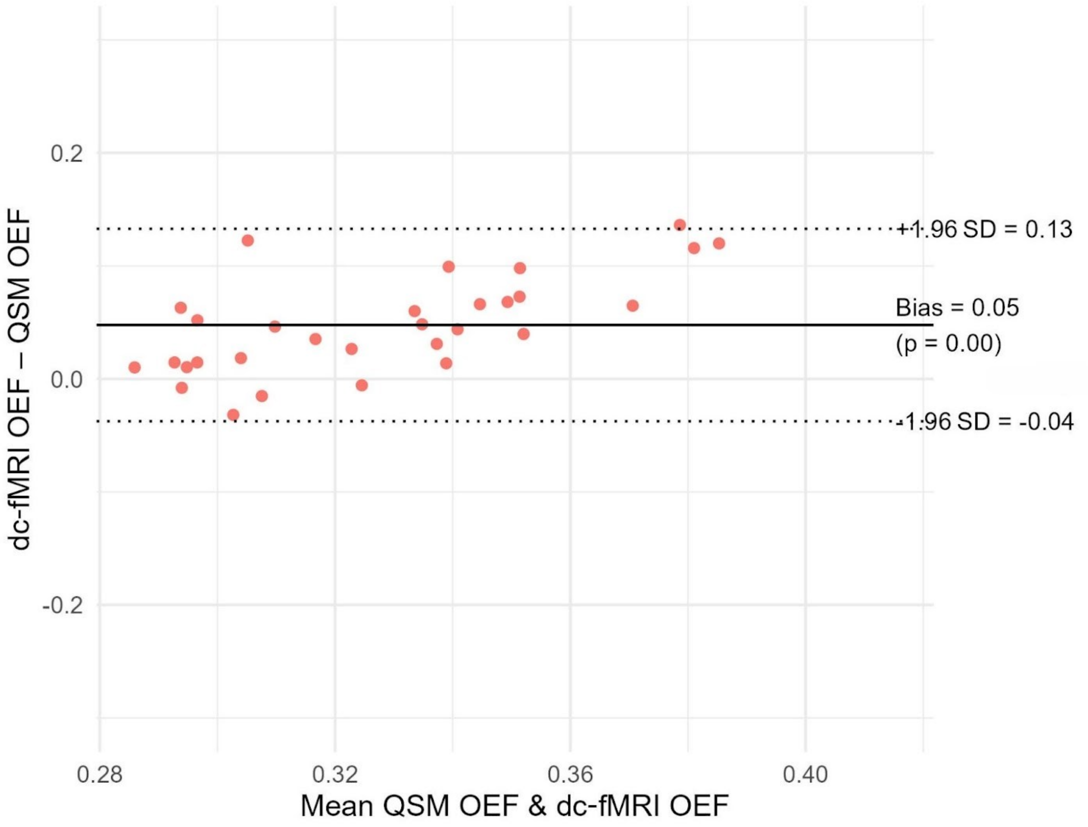


Figure S1. Bland–Altman agreement between GM dual-calibrated BOLD (dc-fMRI) and QSM OEF. Each point represents one participant (x-axis: mean of the two techniques, y-axis: dc-fMRI– QSM). The solid line shows the average bias (+0.05 OEF units), and the dotted lines mark the 95 % limits of agreement (–0.04 to +0.13). QSM systematically underestimates whole brain OEF relative to QSM, yet 95 % of differences lie within the limits and no proportional bias is evident across the observed OEF range.
